# Cingulate cortex shapes early postnatal development of social vocalizations

**DOI:** 10.1101/2024.02.17.580738

**Authors:** Gurueswar Nagarajan, Denis Matrov, Anna C. Pearson, Cecil Yen, Sean P. Bradley, Yogita Chudasama

**Affiliations:** Section on Behavioral Neuroscience, National Institute of Mental Health, National Institutes of Health, Bethesda, MD 20892, USA; NeuroImaging Facility, National Institute of Mental Health, National Institutes of Health, Bethesda, MD 20892, USA; Rodent Behavioral Core National Institute of Mental Health, National Institutes of Health, Bethesda, MD 20892, USA

## Abstract

The social dynamics of vocal behavior has major implications for social development in humans. We asked whether early life damage to the anterior cingulate cortex (ACC), which is closely associated with socioemotional regulation more broadly, impacts the normal development of vocal expression. The common marmoset provides a unique opportunity to study the developmental trajectory of vocal behavior, and to track the consequences of early brain damage on aspects of social vocalizations. We created ACC lesions in neonatal marmosets and compared their pattern of vocalization to that of age-matched controls throughout the first 6 weeks of life. We found that while early life ACC lesions had little influence on the production of vocal calls, developmental changes to the quality of social contact calls and their associated sequential and acoustic characteristics were compromised. These animals made fewer social contact calls, and when they did, they were short, loud and monotonic. We further determined that damage to ACC in infancy results in a permanent alteration in downstream brain areas known to be involved in social vocalizations, such as the amygdala and periaqueductal gray. Namely, in the adult, these structures exhibited diminished GABA-immunoreactivity relative to control animals, likely reflecting disruption of the normal inhibitory balance following ACC deafferentation. Together, these data indicate that the normal development of social vocal behavior depends on the ACC and its interaction with other areas in the vocal network during early life.

## Introduction

Vocal behavior is a critical mediator of social communication through different life stages of many animals, and particularly in social species such as primates^1^. The common marmoset is a small, arboreal monkey with an elaborate repertoire of acoustic calls. While the meaning and usage of most marmoset vocalizations are not well understood, research has shown that different call types convey information about their social organization, environment, and the presence of food or predators^2,1^. Moreover, these calls undergo developmental progression. During the first postnatal months, the acoustic properties and usage of marmoset infant vocalizations change markedly. For example, for different call types, parameters such as duration and frequency follow typical trajectories during the first months of life, transitioning from an immature babbling phase with a mixture of proto-calls to a more discrete and contingent usage of adult-like calls^3,4^. Recent evidence also suggests that parental or social interaction plays a significant role in the proper development of normal vocal behavior, raising the prospect that important aspects of marmoset vocal behavior are learned^5,6^. Thus, a failure to convey appropriate social information through vocal calls could potentially influence how the infant develops its social interactive skills.

In this study, we focus on the anterior cingulate cortex (ACC) and its contribution to vocal behavior and its development in early life. The ACC is a limbic cortical region known to contribute to vocal behaviors^7,8^, and particularly those associated with emotional states^9–11^. Electrical stimulation of the most rostral segment of the ACC elicits vocalizations^12–14^, whereas ACC ablations limit spontaneous vocalizations^15^ and voluntary control of vocal behavior^16,17^. Its dense anatomical connections with the amygdala^18,19^ underscore its role in shaping the affective component of vocalizations^20–23^. At the same time, its descending projections to the periaqueductal gray^24,25^ endow the ACC direct control over activating the brainstem vocalization pathway^7,26,27^.

Vocal production leads to expression of immediate early genes in the ACC^28^, with early studies reporting that infant ACC lesions abolish the characteristic cries that infants normally issue when separated from their mother^29^. These findings implicate the ACC in volitional and emotional control over vocal output.

Longitudinal monitoring of vocal behavior provides a tractable, high dimensional readout of the development of socio-affective circuits. It also provides a means to investigate how early life disruption to brain areas such as the ACC might affect the normal progression of social interaction. If the ACC contributes to the early-life maturation of vocal behavior, then neonatal ACC lesions should hamper the normal control of emotional vocal utterances. Here, we performed excitotoxic ACC lesions in neonatal marmosets and tracked their vocal behaviors, comparing them to age-matched controls throughout the first 6 weeks of life, and examined the impact of the early life lesion on interconnected brain regions in the vocal production network. We demonstrate that animals with neonatal damage to the ACC retained their capacity to issue calls. However, these animals showed a change in their vocal repertoire and an altered acoustic structure in their communicative “social” calls, as well as permanent anatomical changes in the amygdala (AMY) and periaqueductal gray (PAG).

## Results

We studied the vocal behavior in 10 infant marmosets (five males and five females) from five different sets of unrelated parents. In five of the neonatal animals, we performed surgical excitotoxic lesions bilaterally to the rostral portion of the dorsal ACC (24a and 24b) (Fig. 1A, B, see Methods). Starting seven days before the surgery and continuing until six postnatal weeks, infant vocalization behavior was recorded in an isolated, temperature-controlled incubator in 5 min sessions, 2-3 times a week (Fig. 1C, 1D). In four of the animals, the estimate of ACC volume from T2-weighted MR scans performed under anesthesia approximately eight months of age revealed a 60% decrease in ACC volume compared with four control animals (Fig. 1E; two animals were not scanned). Following sexual maturity, at approximately two years of age, the animals were euthanized, and their brains were histologically visualized to verify the extent of the lesion. We also examined downstream effects of the lesion, including an evaluation of its effects on mature neurons (NeuN), inhibitory neurotransmitters (GABA and GAD67), glial cells (GFAP and Iba1) and fiber tracts (myelin; Fig. 1F). See Materials and Methods for details.

**Fig. 1.**
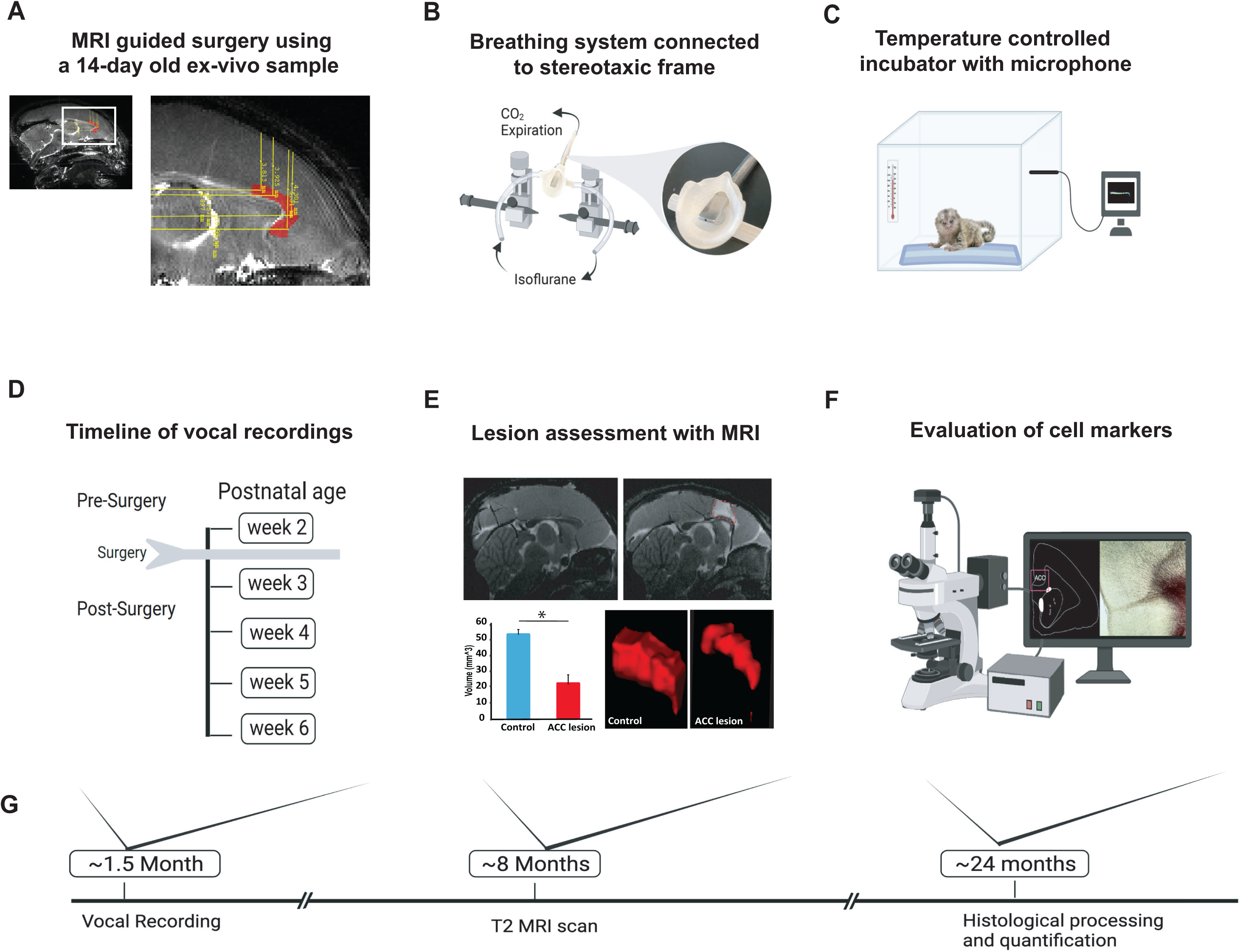
Experimental Design and timeline. **(A)** Reference MR scan was obtained using an ex-vivo sample of a 14-day old marmoset. Parasagittal view of the reference scan shows location of injection coordinates targeting the rostral portion of the dorsal ACC (24a and 24b), bilaterally (red). **(B)** Gas anesthesia was supplied through a custom-made breathing system comprising a facemask fitted with a palate bar with a 0.6mm diameter hole. The palate bar was connected to a vital monitor to accurately detect small tidal end volumes during anesthesia while the animal was secured in the stereotaxic frame. **(C)** Five-minute vocalization recordings were obtained from infant placed in a softly padded temperature-controlled incubator. **(D)** Timeline of vocal recordings from postnatal week 2 to postnatal week 6. The ACC lesion was conducted at postnatal week 2 when animals were 14-16 days old. **(E)** Representative sagittal view of postoperative T2-weighted MR images of a control (left panel) and lesioned (right panel) infant to reveal extent of white hypersignal, which reflects edema due to injections of the excitotoxin and therefore approximate site of the ACC lesion. There was a significant reduction in total ACC volume in the ACC group relative to controls (n=4/per group; F(1,6) = 82.78, p<0.0001). A representative 3-dimensional view of area 24 is presented showing the reduced volume of the ACC (right panel) relative to the normal volume in the control (left panel). **(F)** Schematic illustration highlights the end point of the experiment involving histological processing and evaluation of cell markers. **(G)** Longitudinal timeline shows approximate age of animals following vocal recordings, MRI lesion assessments and histological processing.

### Verification of the ACC lesion and its impact on downstream vocal structures

The intended lesions and reconstructed ACC damage based on histological evaluation are shown in one hemisphere for four animals in Figs 2A and 2B, respectively. The ACC lesion covered most of the target cytoarchitectonic areas 24a and 24b of Paxinos et al.^30^, just above the corpus callosum. The rostral limit of the lesions was adjacent to the genu of the corpus callosum and the caudal limit just anterior of area 23a caudally. Dorsally, the lesions extended past 24b into motor area 6M. There was little if any encroachment into subgenual area 25. Apart from one case which showed some sparing of the lesion in the left hemisphere, there was extensive overlap in the placement of the ACC lesion. The anterior-posterior extent of the bilateral lesion for each monkey is compiled in supplemental Fig. S1 as sections from MRI images.

**Fig. 2.**
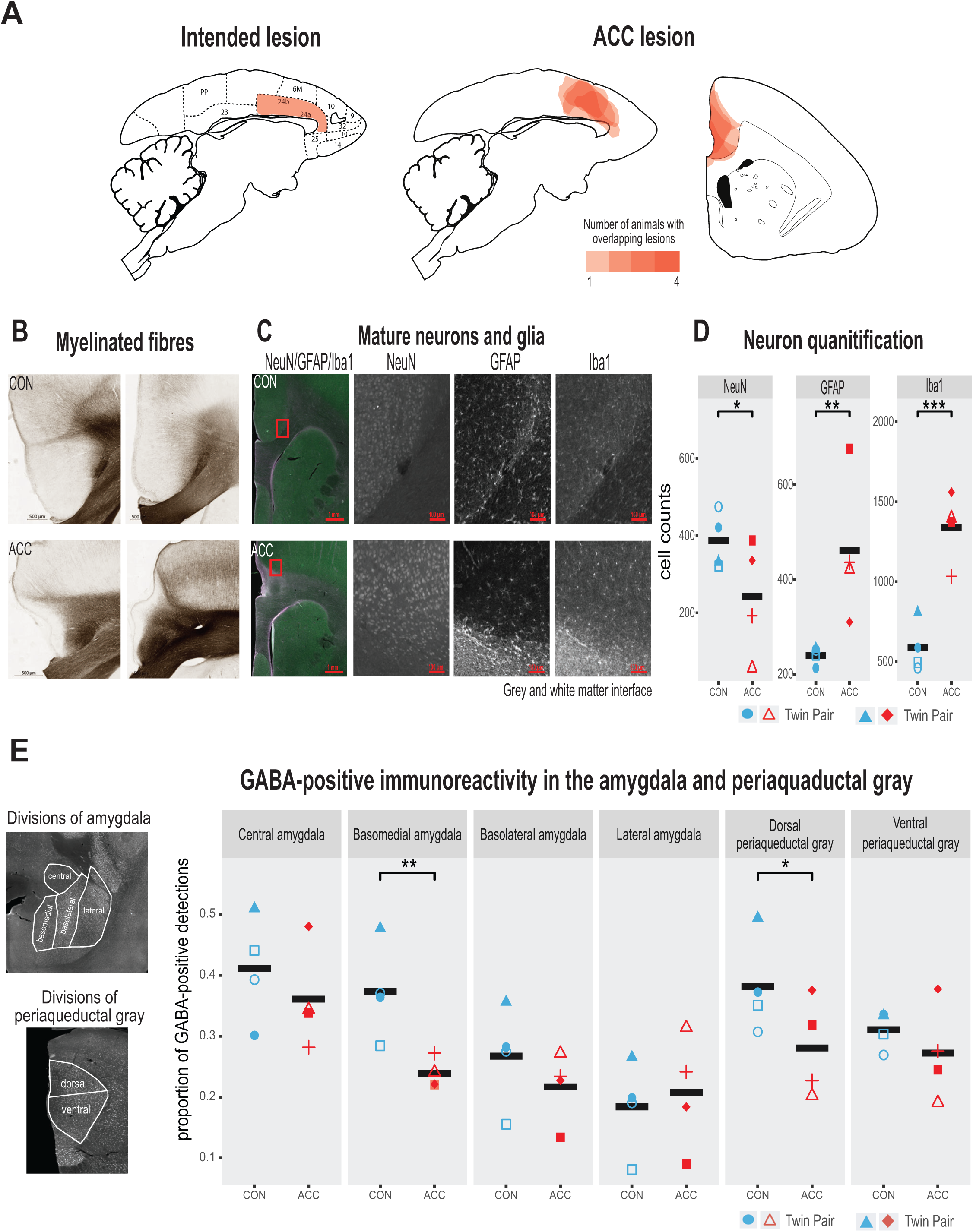
Lesion verification and impact of early life ACC lesion on vocal downstream structures. **(A)** Left panel shows a sagittal section from the standard marmoset brain depicting the intended ACC lesion shaded in red. The right panel shows schematic lesion reconstructions superimposed on a sagittal and coronal marmoset brain section depicting the extent of the ACC lesion shaded in red. Regions that appear darker indicate greater overlap in the damage present among different animals. **(B)** Magnified images of area 24 stained to visualize myelinated fibers in a representative control (top panel) and ACC-lesioned (bottom panel) animal. The normal radial arrangement of the myelinated fibers is disrupted following the ACC lesion. **(C)** Histological visualization of mature neurons and glia in representative control (top row) and ACC-lesioned animal (bottom row). Leftmost image shows neurons (green), astrocytes (violet) and microglia/macrophages (white) in the same image. Red square represents the magnified grayscale sections showing cell loss (NeuN), high levels of astrocytes (GFAP) and microglia/macrophages (Iba1) surrounding the lesion site at the grey and white matter interface in the ACC-lesioned animal (bottom row) relative to the controls (top row). **(D**). Histological quantification of mature neurons and glia in cortical area 24 in the ACC-lesioned animals, or in the corresponding intact cortical tissue bordering white matter in the controls. There was a reduction in the number of mature neurons (NeuN) and an increase in glia (GFAP and Iba1) in the ACC group relative to controls. *p<0.05; ** p<0.001; *** p<0.00001. **(E)** Left panel shows grayscale images with anti-NeuN staining depicting divisions of AMY and PAG where relative distribution of GABA-positive immunoreactive expression were quantified. Right graphs show the proportion of GABA expression in each division depicted in the AMY and PAG. Each circle represents one animal (ACC lesioned animal is red, Control animal is blue). Mean expression is represented by black bar. Data for two animals in ACC group overlap for basomedial AMY quantification. GABA-positive immunoreactivity was significantly down in the basomedial AMY and dorsal PAG.

Representative photomicrographs of the ACC lesion and a control are presented in Fig. 2B, which shows the distribution of myelinated fibers in the ACC region stained using a high-resolution Black-Gold II myelin stain (Histo-Chem Inc., Jefferson, AR). There was clear evidence that the lesion created a major disruption to the normal radial arrangement of the fibers in the ACC region caused by extensive demyelination of the axons (Fig 2B). The ACC lesion also impacted the integrity of white matter tracts local to the site of the lesion (Fig. S2), but the transverse diameter of major fiber tracts, namely the corpus callosum and the anterior commissure did not differ between the groups. The loss of neurons, however, and the respective increase in glial cells at the lesioned site especially at the interface between the gray and white matter was clearly observed (Fig. 2C, D).

We examined the cellular and neurotransmitter composition of the amygdala (AMY) and periaqueductal gray (PAG), as these structures are downstream from the ACC and their natural development may be affected by the infant ACC lesions. We first investigated whether neurons in these structures were degenerated using Fluro-Jade C (Histo-Chem Inc.), which is used as a marker for apoptotic, necrotic, and autophagic cells. There was no sign of neurodegeneration in these downstream brain regions two years following the infant lesion. We next examined the proportion of neurons in the AMY and PAG expressing GABA, since changes in the relative number of inhibitory interneurons could serve as a marker for downstream neuroplasticity in response to the ACC lesion^31^. We found a significant reduction of GABA positive neurons in two structures, the basomedial AMY and the dorsal portion of the PAG (Fig 2E). This reduction suggests a disruption of the normal inhibitory balance within the vocal network following the infant ACC lesions.

### Vocal behavior persists immediately following neonatal ACC lesions

In the weeks following bilateral ACC lesions, infants tested in the isolated chamber continued to vocalize readily, which is somewhat surprising given the critical role of this structure in normal vocal behavior^12–14^. From (presurgical) postnatal week 2 to (postsurgical) postnatal week 6, we annotated 23,000 calls from the five lesioned and five control marmosets. Sample spectrograms from audio recordings of a twin pair before and after surgery are shown in Fig. 3A-B. While there was variability among individuals, the calls were complex and diversified from postnatal week 2, consisting of cries as well as immature versions of adult vocalizations including phee, twitter, and trills, as well as complex calls when two calls merged such as trill-twitter or cry-phee, or any other combination (Fig 3A). By the sixth postnatal week, the call repertoire for both the lesioned animals and the controls had both evolved, with no conspicuous difference between groups (Fig 3B). Most notably, the relative reduction of the total rate and diversity of calls was similar between groups (Fig. 3C, χ2(2)=2.8464, p=0.24), with the reduction matching the known maturational changes accompanying growth of the vocal apparatus and increased respiratory powers^32^. Further analysis, on the proportion of calls emitted following surgery showed that the ACC lesion had minimal effects on the rate of most call types during this period (phee (β= −0.07, 95% CI [-0.31, 0.17], p = 0.49); twitter (β= −0.07, 95% CI [-0.16, 0.01], p = 0.09); trill (β= −0.03, 95% CI [-0.11, 0.06], p = 0.49); cry (β= 0.13, 95% CI [-0.03, 0.29], p= 0.10) (Fig 3D). The exception to this rule was an elevation in the rate of ‘other’ calls, which comprised tsik, egg, ock, chatter and seep calls. These calls were significantly elevated in animals after the ACC lesion (β= 0.11, 95% CI [0.03, 0.20], p = 0.018). This was driven mostly by an increase in the lesion group during postnatal week 4.

**Fig 3.**
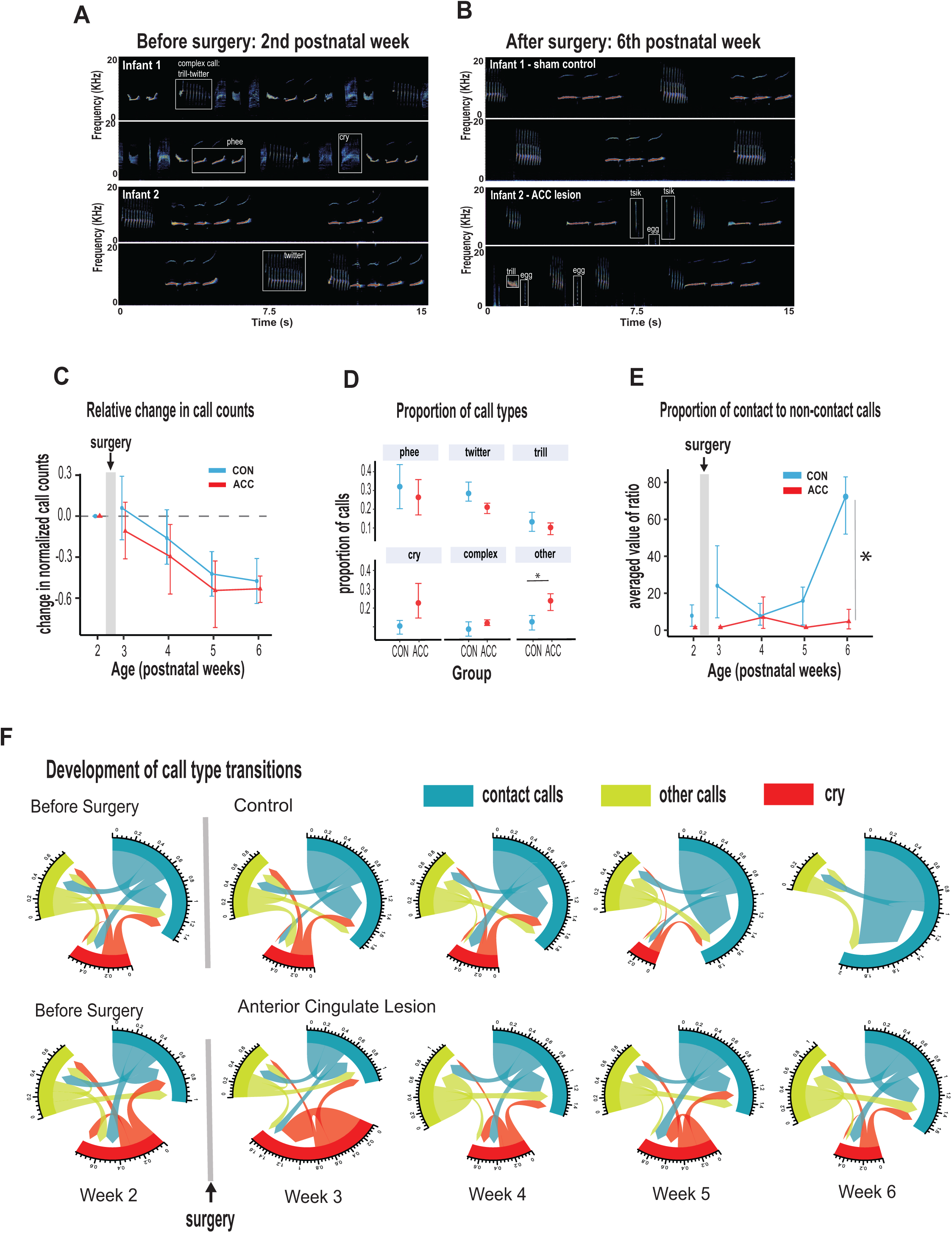
ACC in early life is integral to postnatal development of social contact calls. **(A-B)** Spectrograms show sample 30-second vocal recordings of a representative control and ACC-lesioned marmosets before (postnatal week 2) and after surgery (postnatal week 6). Before surgery, the infants ‘babbled’ by emitting a wide range of immature concatenated calls, each with its own spectrogram motif illustrated and labeled in boxes. After surgery, at postnatal week 6, calls show reduced variability separated by distinct gaps or inter-call intervals. **(C)** Both groups show a reduction in the relative call count with increasing age. **(D)** Average proportion of each call type pooled from week 3 to week 6 following surgery. Animals in both groups were able to emit calls of different call types. Those with ACC lesions made minor calls designated as ‘other’ more frequently than controls but all major call types were produced at equivalent rates. (E) Proportion of social contact calls relative to non-social contact calls. The y-axis represents the averaged value of the ratios of the number of social calls divided by the number of non-social calls (*x (# social calls / # non-social calls).* Despite their ability to produce all call types, the proportion of social contact calls comprising phee, twitter and trills, was substantially reduced in animals with early life ACC lesions at postnatal week 6. Due to factors beyond our control (Covid-19), the number of recordings varied between animals: week 3: CON n=5, ACC n=5; week 4: CON n=5, ACC n=4; week 5: CON n=4, ACC n=3; week 6: CON n=4, ACC n=3. **(F)** Chord diagrams illustrate the likelihood of transitioning between call types. At 6 weeks of age, animals with ACC lesions showed a higher likelihood of transitioning between all call types, but less frequent transitions between social contact calls relative to the sham group. The chord diagrams visualize the weighted probabilities and directionality of these transitions between different call types. Weighted probabilities were used to account for variations in call counts. The thickness of the arrows or links indicates the probability of a call transition and the numbers surrounding each chord diagram represent the relative probability value for each specific transition.

Two additional variables relatively unaffected by the ACC lesion were the call durations and inter-call intervals, acoustic features that have been used to track vocal development in previous studies^4,6,33^. Consistent with the overall decrease in vocalization rate with increasing age, there was an associated increase in inter-call intervals which was noted at late postnatal weeks (β= 0.33, 95% CI [0.22, 0.43], p < .001) which held true for both controls and lesioned group (χ2(2)=1.88, p=0.39). None of the specific call types exhibited developmental changes in call type duration (phee (χ2(2)=1.08, p=0.58); trill (χ2(2)=2.87, p=0.24); twitter (χ2(2)=2.79, p=0.25); cry (χ2(2)=0.057, p=0.97), but there were slight changes in the duration of phee syllables exhibited only by animals with an ACC lesion which we discuss later. Importantly, we did not observe ACC-lesion induced changes in physical growth factors such as body weight and grip strength which could feasibly impact vocal parameters such as duration^4^ (Fig. S3).

### Neonatal ACC lesions prevent early maturation of social contact calls

Whereas many of the basic vocal parameters evolved normally in the animals with the ACC lesions, one major difference related to their use of social contact calls. By 6 weeks of age, marmoset vocalizations are known to approach their mature state and become dominated by social contact calls, namely phees, trills, and twitters. The phee call is studied most extensively as a long-distance contact call. It is typically evoked when the animal is socially distanced or isolated, and it promotes vocal exchanges between marmosets located out of sight in far-away locations^34,35^ to facilitate reunion with family groups^36^. Trills and twitters are short distance contact calls thought to monitor the presence of group members^37,38^. Since the ACC plays a major role in socioemotional cognition (for review, see Devinsky et al.^39^), we surmised that the ACC lesion might specifically influence the socioaffective content of the vocalization that is normally expressed through contact calls.

We thus grouped phee, twitter, and trill calls as social contact calls and compared them with non-contact calls (i.e., cries and other minor occurring calls like tsik, egg, ocks, chatter, and seep). At 6 weeks of age, social contact calls predominated the control animal’s vocalization. However, in ACC-lesioned animals, this aspect of social vocal behavior was substantially reduced. This difference emerged gradually after the surgery, and was only evident at 6 weeks of age (Fig 3E; (χ2(2)=7.58, p=0.022). By the sixth week, the social vocal repertoire of the lesioned animals was altogether different from the control animals, with a much smaller proportion of social contact calls.

To further understand the effect of the ACC lesion on the normal distribution of calls, we investigated the call transition probabilities between contact calls, cries, and other calls (Fig 3F). In contrast to the control animals, whose repertoire was dominated by social contact calls, the ACC lesion group showed frequent transitions mostly to other non-contact call types (u-index Wilcoxon test, p = 0.055). These data suggest, therefore, that the ACC mediates developmental changes within the first 6 weeks of life that lead to the dominant production of isolation-induced contact calls and the gradual reduction of cries and other calls.

### Neonatal ACC lesions alter sequential characteristics of social contact calls

We examined the characteristics of vocal sequences to learn more about how early life ACC lesions might influence the acoustic signals that marmosets potentially relay to distantly located family members or other conspecifics when socially isolated. We focused on phee syllables, which are discrete elements or components of a call separated by very short intervals^39,40^. Thus, a sequence of phee calls may comprise multiple syllables (Fig 4A). The functional significance of syllables is not clearly understood but a change in the number of syllables or their acoustic characteristics might feasibly alter the message conveyed to a family that cannot be seen or heard. This is especially important to young infant monkeys that are naturally demanding of attention, and even more so if isolated. The number of syllables per phee call was highly variable amongst the animals, ranging from 1 to 8 syllables per phee.

**Fig. 4.**
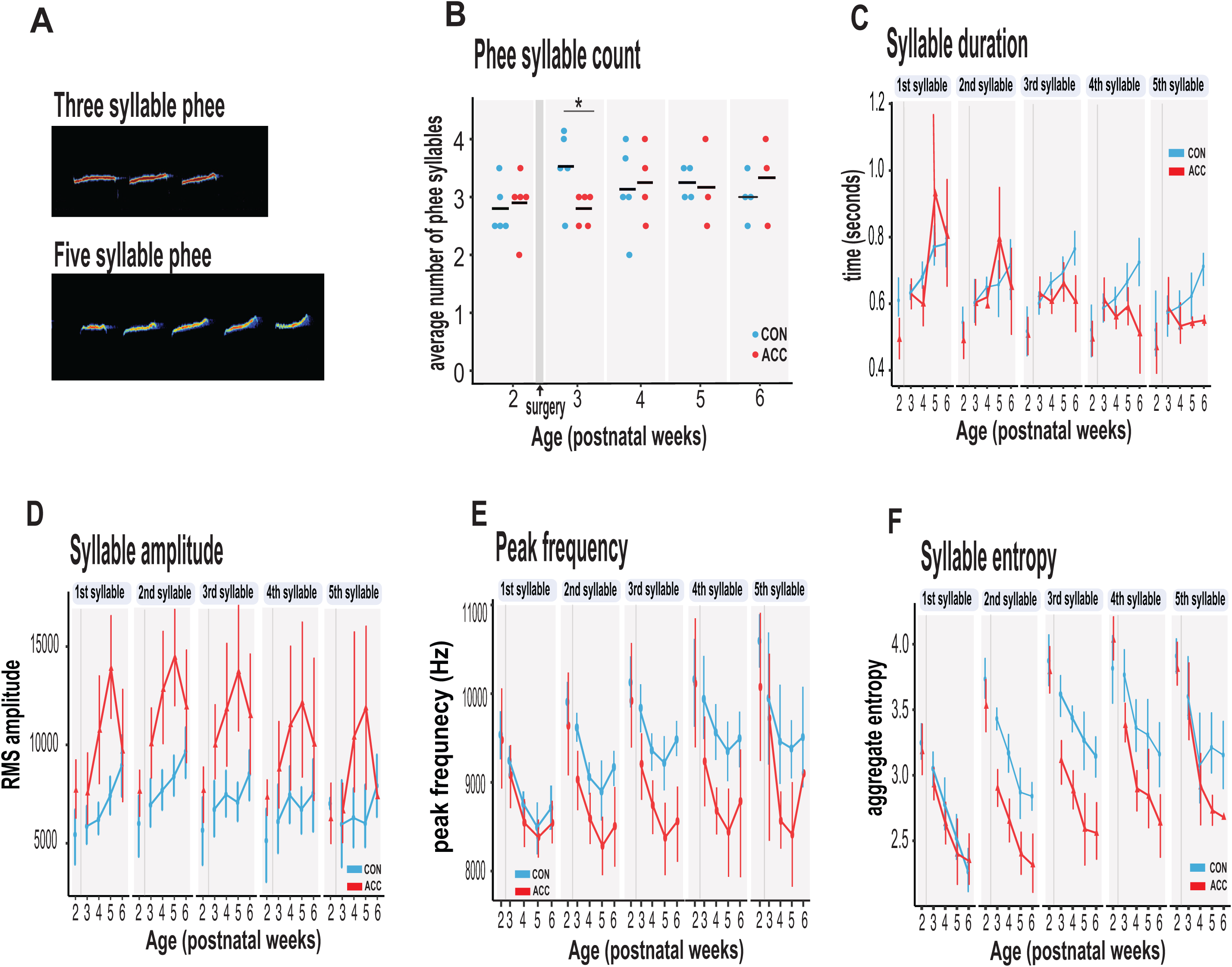
ACC lesion alters structural characteristics of long distance social phee calls. **(A)** Sample spectrogram with examples of three and five syllable phee calls **(B)** The ACC lesion caused a reduction in average phee syllable counts immediately after the ACC lesion (red dots) at postnatal week 3, but then normalized to 3-4 syllables thereafter. **(C)** Phee syllable duration ACC-lesioned group became shorter for multisyllabic phees ≥ 3 especially with increasing age. **(D)** The effective amplitude for each phee syllable increased for the ACC group until postnatal week 6. **(E)** Animals with ACC lesions emitted low entropy phees for calls as low as 2-syllables and continued until postnatal week 6**. (F)** Decrease in peak frequency of phee calls immediately following ACC lesion. Data in D-F represents average of specific syllable in a phee sequence irrespective of the number of syllables in a phee (e.g., 1 syllable phee, the 1st syllable in a 2-syllable phee, in a 3- syllable phee and in a 4-syllable phee). Due to factors beyond our control (Covid-19), the number of recordings varied between animals: week 3: CON n=5, ACC n=5; week 4: CON n=5, ACC n=4; week 5: CON n=4, ACC n=3; week 6: CON n=4, ACC n=3. Error bars are confidence intervals. Gray shaded lines or bars represent time of surgery.

The ACC lesion did not greatly affect the phee syllable count. Aside from a transient decline in the number of syllables in the week after the surgery (postnatal week 3: Wilcoxon test p=0.042), these animals showed the normal preferred range of 3-4 phee syllables at later postnatal weeks (Fig 4B). Even in the sixth postnatal week, when the proportion of phee and other contact calls was much lower in the ACC lesioned animals, the number of syllables in those phee calls that were issued was similar to the control group.

However, other phee call variables were affected by the lesion. For example, the duration of phee syllables was shortened in ACC-lesioned animals (χ2(5)=13.27, p=0.021), particularly in the later syllables of a multi-syllablic phee. This effect emerged gradually and was most pronounced when the animals were 5-6 postnatal weeks (Fig. 4C). Likewise, the amplitude of phee syllables was also affected by the ACC lesion (χ2(5) = 48.178, p<0.0001) with lesioned animals making louder phee calls (Fig 4D). For each phee syllable, the amplitude difference between groups increased until postnatal week 5 and then disappeared at postnatal week 6 (postnatal week 4: β = 3017.75, 95% CI [1831.61, 4203.89] p < .001; postnatal week 5: β = 3719.00, 95% CI [2389.88, 5048.12], p < .001). In line with the increase in amplitude, the peak frequency of each phee syllable was also lowered by the ACC lesion (Fig 4E). This change occurred soon after the lesion [postnatal week 3: β = −488.63, 95% CI [−702.40, −274.85] p < .001; postnatal week 4: β = −384.61, 95% CI [−607.98, −161.24], t(513) = −3.38, p < .001].

Finally, we examined entropy of phee syllables as a measure of vocal complexity^72^. High entropy in multisyllabic phees would indicate that these vocalizations are diverse, variable and unpredictable. We found that animals with ACC lesions exhibited lower entropy in phee syllables relative to controls as early as postnatal week 3 (χ2(5) = 34.528, p < .0001) (Fig 4E) suggesting that their vocal sequences were simple, less diverse and stereotyped. Together, our data suggest that the ACC lesion compromised the normal development of the phee signature for each monkey by making them shorter, louder, and monotonic.

## Discussion

We found that early life ACC lesions led to rather specific alterations in the production of vocal calls, developmental changes to the quality of social contact calls, and their associated sequential and acoustic characteristics were compromised. Contact calls that normally dominate the marmoset vocal repertoire at around 6 weeks were selectively diminished in the lesioned animals. When one common contact call, the phee call, was issued, its structural characteristics were unusual. The ACC lesions also led to permanent changes in remote brain areas known to be involved in vocal behavior. Notably, the proportion of presumptive inhibitory interneurons was reduced in both the basomedial AMY and dorsal PAG. We can infer from these findings that an intact ACC in early life is integral to postnatal development of social vocalizations, and that its interactions with vocalization-eliciting sites from a very early age is fundamental to the normal vocal expression of social behavior.

Consistent with previous reports, the range of call types observed in both neonatal controls and neonatal ACC-lesioned animals within the first weeks of postnatal development were stereotyped and repetitive^33,41^. With increasing age, the call rate gradually declined such that by the time the animals were six weeks of age, the most common vocalizations were those that conveyed social distance. Such calls solicit attention from family members and may trigger a range of behaviors including search, approach, interaction, and caregiving in response to the need for social contact. This is especially so for infants whose well-being depends on social feedback and reciprocal interaction^4,5,35,42^. Our data suggest that the ability to effectively convey this social need was significantly altered in animals with early life ACC damage. At 6 weeks of age, these animals were not making social vocalizations at the same high rate as their age-matched controls. This reduction in social vocalizations does not appear to reflect a general slowing in vocal maturation, since other call types had advanced at the normal rate.

Since ACC lesions in humans cause social apathy^43,44^, one possibility is that early life ACC removal altered the animals desire or motivation for social reinforcement; these infants appeared to make little effort in using vocalizations to solicit social contact when socially isolated. This change in call usage aligns with their social development period at around postnatal week 6 when infant marmosets transition from using fixed, stereotypical calls to a flexible and more individualized call repertoire as they wean towards independence^33^. Our data suggest that this transition does not occur normally following an ACC lesion.

In addition, the neonatal ACC lesion altered the quality of the infants’ long-distance phee calls; they were shorter in duration, louder in amplitude, lower in peak frequency and abnormal in their entropy such that the acoustics of calls were blunted of variation and less diverse. This suggests that the social message conveyed by these infants to their families through phee calls, even though it was loud and propagating over long distances, was potentially deficient, limited, and/or indiscriminate. However, the impact of entropy on emotional quality of vocalizations has not been systematically explored. Generally speaking, high entropy relates to high randomness and distortion in a signal. Accordingly, one view posits low-entropy phee calls represent mature sounding calls relative to noisy and immature high-entropy calls^42^. In the current study, the reduction in syllable entropy observed for both groups of animals with increasing age is consistent with this view.

At the same time entropy relates to vocal complexity; high entropy refers to complex and variable sound patterns whereas low entropy sounds are predictable, less diverse and simple vocal sequences^72^. One possibility is that call maturity does not equate directly to emotional quality. In other words, a low-entropy mature call can also be lacking in emotion as observed in humans with ACC damage; these patients show mature speech, but they lack the variations in rhythms, patterns and intonation (i.e., prosody) that would normally convey emotional salience and meaning. Our observation of a reduction in phee syllable entropy in the ACC group in the context of being short, loud with reduced peak frequency is consistent with this view and suggests that the ACC group were emitting phee calls that were potentially lacking emotional meaning.

The long-term behavioral implications of such imperfect vocalizations is currently unknown but could, ostensibly, affect their ability to use long distance social vocalizations to maintain intragroup functions such as warn of predators, strengthen family bonds, and maintain group cohesion more generally^2^. Since the ACC exerts regulation over autonomic responses^45–47^, its ablation so early in life might feasibly blunt respiratory and vocalization responses in negative emotional environments such as social isolation. How these factors impact vocal behavior is a current topic of investigation (Sheikhbahaei et al., *SfN* abstracts. 2023).

Although we found that an intact ACC in early life is integral to the postnatal maturation of social vocalizations, we also show that it is not critical for production of infant vocalizations more generally. A number of early observations reporting the loss of learned or spontaneous vocalizations following bilateral ACC lesions, left this question open, though it has been clear that vocal production in adults can withstand ACC damage^16,17,48–50^. In addition to ablating the ACC, researchers found that it was necessary to ablate other frontolimbic areas to permanently eliminate infant vocalizations such as cries^17,29^. Our findings indicate that innate vocal production in the earliest phases of life, as early as two weeks postnatally in the marmoset, is not critically dependent on the ACC. The infant ‘babbling’ behavior observed in marmosets and other primates^4,41,51^ was largely preserved in the ACC-lesioned infant monkeys which, like the control group, produced long sequences of concatenated calls composed of rudimentary features of mature adult-like calls.

While the ACC is not essential for infant vocal behavior, its absence affects not only the maturation of social vocal behavior, but also the anatomical compositions of structures with which it is interconnected. We noted a decrease in presumptive inhibitory interneurons in the dorsal PAG and basomedial AMY, two prominent ACC target regions involved in vocal behavior^7,20,52^. We can speculate that this reduction might stem from a prolonged deafferentation of cingulate inputs, gradually leading to a rebalancing of the excitatory/inhibitory elements in the local circuit.

One potentially related observation is that phee calls became louder in the weeks following the surgery. It is interesting to speculate that such amplitude increases might reflect a local decrease in inhibition in structures such as the AMY or PAG, whose activity is thought to tune the emotional characteristics of social vocalizations. The primate ACC receives dense projections from the basomedial AMY with notably minimal direct input from the lateral nucleus^53–55^, and layer V pyramidal neurons project to the PAG^56,57^ with a greater concentration directed to the dorsolateral column^58^. Thus, the ACC has the capacity to directly activate the brainstem vocalization pathway as early as the first few weeks of life. Both, the AMY and PAG are highly active during contexts in which threat related vocalizations would normally be triggered^59–61^, and both regions elicit vocalizations through electrical stimulation or pharmacological disinhibition^12,62–64^. We cannot be sure when during postnatal development the ACC lesion altered GABA expression in the AMY and PAG, but from our results, we can infer that the appropriate modulation and coordination of social vocal behavior requires the normal postnatal development of the ACC.

Existing evidence in monkeys and humans demonstrate unequivocally the importance of the ACC in its contribution to emotional vocalization. In humans, as in monkeys, ACC lesions do not eliminate vocal behavior, but instead tend to remove the intonation and prosodic features of the vocalization characterized as expressionless^49^. This is consistent with the changes observed in the marmoset phee calls. In general, our data suggest that the ACC shapes the emotional structuring of social calls during the first few weeks of life in the marmoset. Given the many similarities to humans, and the strong contribution of socioemotional information to the vocal productions beginning in infancy, it is reasonable to speculate that similar mechanisms apply to the development of early life human vocal behavior. Importantly, our data confirm the importance of vocalizations as a means of conveying social information even when family members or conspecifics are not physically present. Our data suggest that in the absence of a functioning ACC in early life, infant calls conveying social information that would elicit feedback from parents and other family members may be compromised, and this could potentially influence how that infant develops its social interactive skills. The ability to normalize brain circuits in early life would provide a major therapeutic advance for the remedial treatment of social deficits that plague disorders of mental health.

## Materials and Methods

### Subjects

All procedures accorded with the Guide for the Care and Use of Laboratory Animals and were approved by the Animal Care and Use Committee of the National Institute of Mental Health. A total of 10 marmosets (Callithrix jacchus), 5 males and 5 females, all born in captivity, were used in this study. Five infants received ACC lesions at 14-16 postnatal days old. Five others served as age-matched controls. The infants were raised by parents and siblings in family groups comprising 4-6 members and housed in temperature-controlled rooms (∼27°C), 50-60% relative humidity under diurnal conditions (12h light:12h dark). Food and water were available ad libitum, supplemented with fresh fruit or vegetables. One animal showed sparing of the ACC lesion in one hemisphere. The final sample size for the behavioral data was n=5/per group. For the MRI and histological data, the final sample size was n=4/per group.

### Surgery

We first obtained a reference MRI scan using a 14-day old ex-vivo sample (Fig. 1A). A T2 weighted scan was obtained using 7T Bruker Biospin MRI platform with an eight-channel volume coil. Using ParaVision Acquisition 6.0.1, the following echo sequence was used to acquire a 3-dimensional volume of the infant marmoset brain: TR = 400, TE = 72ms, flip angle= 90 degrees, matrix size = 256x256x214, resolution = 0.15 mm isotropic, number of averages = 8, number of repetitions =1) and the total scan time was 3 hrs. The scan was aligned horizontally by rotating the image until the anterior and posterior commissures were positioned at the same height and water filled ear bars were used to obtain the interaural reference. We then used ITK-SNAP^65^ (Yushkevich et al., 2006) to identify the anterior cingulate cortex (ACC) at the coronal planes before the genu of corpus callosum to the level of anterior commissure. The coordinates were calculated relative to the ear bars and midline references, both of which were visible on the scan. The resulting 14-day old marmoset scan served as a template atlas to calculate injection coordinates to target the rostral portion of the dorsal ACC (24a and 24b), bilaterally, in all marmosets. We calculated 5 injection coordinates for each hemisphere: (1) AP: 10.7 mm, ML: ±0.7 mm, DV: -2.9 mm; (2) AP: 10.7 mm, ML: ±0.7 mm, DV -4.5 mm; (3) AP: 9.5 mm, ML ±0.7 mm, DV −3.5 mm; (4) AP 8.5 mm, ML ±0.7 mm, DV −3.6 mm; (5) AP 7.5 mm, ML ±0.7 mm, DV −3.5 mm.

The entire surgical procedure was performed under aseptic conditions in infant monkeys that were 14-16 days old. During surgery, monkeys received isotonic fluids. Heart and respiration rates, body temperature, blood pressure, and expired CO2 were monitored throughout the procedure. Pre- and post-operatively, monkeys received non-steroidal anti-inflammatory drugs, (meloxicam, 2mg/ml, s.c) to reduce swelling. The monkey was first immobilized with an anesthetic dose of alfaxalone (10mg/kg, i.m) combined with diazepam (5mg/kg, i.m). In this state, the infant’s head was shaved, and vital electrodes were secured on the infant’s chest. Temperature was measured with a rectal probe. The infant’s head was then secured in a small animal stereotaxic frame (Stoelting Company, Illinois) attached to a custom-built stage fitted with eye bars, ear bars and a pallet bar to accommodate the small head. Once the head was secured in the frame, anesthesia with isoflurane gas (1-2% to effect) was provided through the custom fitted mask (Fig. 1B). An integral part of the pallet bar was a gas hole (0.6 mm diameter) that ran along the length of the bar and connected, via tubing, to a vital monitor to measure small end tidal CO2 volumes.

Following a midsagittal incision, the scalp was retracted, and a craniotomy was made above the target coordinates of the brain. A 5 µl syringe (33 gauge, Neurosyringe, Hamilton Company, Reno, NV) was used to administer bilateral injections of 0.12M NMDA (M3262, Sigma-Aldrich) dissolved in sterile filtered saline into the anterior cingulate cortex (0.5 µl per injection site). Each injection was made over 2 mins and the injector remained in place for an additional 4 mins for dispersion. When all injections were complete, the scalp was closed with intradermal absorbable sutures and the infant was allowed to recover in an intensive care unit that was void of extraneous sensory stimulation (e.g., excessive bright lights and loud noise). During recovery, marmosets received a combination of Esbalic and Enfamil (3:1 ratio) infant formula every 2-3 hours. When fully awake, each infant was returned to its family unit. A total of five marmosets received the neonatal cingulate lesion. Another five marmosets served as controls: two received saline injections (shams), one received a craniotomy only, and another two were unoperated.

### Vocal recordings

Each infant was placed in a temperature-controlled incubator set to ∼38°C (Thermocare, CA), and the emitted vocalization was recorded for 5 mins. Sound recordings were acquired using a cardioid microphone (Sennheiser ME 64, Sennheiser, Wedemark, Hanover, Germany) that was placed on the side of the incubator (Fig. 1C). The microphone was connected to a computer and recorded sounds were digitized at a sampling frequency of 44 kHz using Raven Lite software (Cornell Lab of Ornithology, Ithaca, NY). Due to a variety of extraneous factors beyond our control including restrictions due to the COVID-19 pandemic, the exact day and number of recordings differed between monkeys. Therefore, recording sessions from each infant were grouped by week. All recording sessions were conducted without the presence of investigators in the recording room. The infant was then returned to its family unit.

### Acoustic Analysis

The spectrogram of each audio file was obtained and visually inspected using Raven Pro 1.6 (Cornell Lab of Ornithology, Ithaca, NY). Spectrogram were generated with a Hann window of 512-sample points to filter the signal at 3dB bandwidth of 124 Hz (example, Fig. 2 A-B). The calls were manually classified by a defined classification system^37,51^. To identify call types in the spectrogram of a recording, Raven software features, such as amplitude waveform, spectrogram and audio playbacks were used. Six major call types (phee, trill, trill-phee, twitter, cry, and complex calls) were identified from spectrograms, along with other minor call types (tsiks, chatter, egg, ock, seep). In some cases, when trill-phees and phees looked similar in spectrograms of a recording, acoustic parameters such as entropy were used to carefully classify calls. Complex calls, comprised vocalizations with elements from at least two different simple call types such as trill-twitter or twitter-phee, etc. From each recording, call types were manually annotated by 3 trained investigators (inter-rater reliability >80%). The spectrograms were used to obtain acoustic measurements such as peak frequency, RMS-amplitude and aggregated entropy and exported for further analysis.

Acoustic characteristics such as inter-call interval, syllable duration, number of phee syllables were quantified using a custom-written R script. Acoustic analyses were performed only for phees, which served as the major call type because of their abundance during postnatal weeks. In some cases, vocalization quantity and amplitude was largely suppressed for several minutes after handling by the experimenters. Consequently, analysis of each recording began after 2 minutes had elapsed.

### Lesion Assessment with MRI

Gross ACC volume was measured from anesthetized MR scans performed at approximately 8 months of age. Anesthesia was induced with 5% isoflurane. The animals were then placed in an MR compatible cradle where its head was secured using ear bars. Isoflurane was maintained at 1.5 - 2.5% and vitals were monitored with V9004 Series Capnograph Monitor (San Clemente, CA). We obtained T2 weighted scan (n=4 Control and n=4 for ACC) using the MR procedure described above with the following echo sequence: TR = 30, TE = 48, matrix size = 144x144x128, resolution = 0.25 mm isotropic, number of averages = 8, number of repetitions =1. The rostro-caudal extent of the ACC was segmented and measured using ITK-SNAP 4.0. A representative sagittal view of the lesioned and non-lesioned ACC can be seen in Fig. 1E. Voxels containing ACC were carefully labeled from the anterior to posterior slice of the MR scan for each subject.

### Histological preparation and quantification

At approximately 24 months of age, the marmosets were euthanized and perfused with 0.1 M PBS followed by 4% paraformaldehyde. The brains were extracted and cryoprotected in 0.1 M phosphate buffered sucrose (in steps of 10%, 20%, and 30% w/v). The brains were partitioned along the midline and right hemispheres were used for further histological processing after sectioning in coronal orientation on a sliding microtome and cryostat into 40 μm sections.

### Immunohistochemistry

The immunohistochemistry (IHC) was performed on two series of free-floating sections. Initially, from each of the 8 brains, six sections each were collected around three rostrocaudal planes at the following approximate locations (in reference to the interaural axis): +13.30 mm (target brain region: the anterior cingulate cortex); +9.20 mm (target brain region: the amygdala); and +2.05 mm (target brain region: the periaqueductal gray matter). Subsequently, these 18 sections were divided into two IHC series of 9 sections each, where three sections covered each target brain area, for separate processing. The first IHC series was processed to visualize major cell classes that are likely to be affected by a brain lesion (neurons: primary antibody against NeuN, astrocytes: primary antibody against the glial fibrillary acidic protein (GFAP), microglia/macrophages: primary antibody against the ionized calcium-binding adaptor molecule 1 (Iba1). The second IHC series was processed to visualize and assess the ratio of GABA- ergic neurons to all neurons: such changes in the relative number of inhibitory neurons could indicate local downstream neuroplasticity in reaction to the ACC lesion, as GABA-ergic neurons are primarily responsible for local inhibition. In the second IHC series we visualized the distribution of NeuN, as well as of neurotransmitter GABA and GAD67, a rate-limiting enzyme in GABA synthesis that produces more than 90% of GABA in the central nervous system.

The first IHC series was incubated in a cocktail of the following primary antibodies for 60 h at 4℃: NeuN (chicken, ABN91, Millipore Sigma), 1:500 dilution; GFAP (goat, SAB2500462, Millipore Sigma), 1:1,500 dilution; Iba1 (rabbit, 019-19741, Fujifilm Wako Chemicals), 1:1,500 dilution). Then after several washes the sections were incubated in a cocktail of the following secondary antibodies for 2 h at RT: 1:200 donkey anti-chicken Alexa Fluor 488 (703-545-155, Jackson ImmunoResearch), 1:500 donkey anti-goat Alexa Fluor 594 (A21207, Invitrogen), 1:400 donkey anti-rabbit Alexa Fluor 680 (711-625-152, Jackson ImmunoResearch). Additional antibodies used in the second IHC series were as follows. Primary: GABA (rabbit, A2052, Millipore Sigma), 1:600 dilution; GAD67 (mouse, MAB5406, Millipore Sigma), 1:500 dilution. Secondary: 1:500 donkey anti-rabbit Alexa Fluor 594 (A21207, Invitrogen), 1:400 donkey anti-mouse Alexa Fluor 680 (A10038, Invitrogen). Finally, the sections were mounted on a glass slide, air-dried, and cover-slipped with DEPEX mounting media (13515, Electron Microscopy Sciences, Hatfield, PA).

### Myelin staining

Myelinated fibers were stained with aurohalophosphate-based Black-Gold II (Histo-Chem Inc., Jefferson, AR) as shown in Schmued et al.^66^ according to the manufacturer’s protocol and cover-slipped with Permount (SP15, Fisher Scientific).

### Staining for degenerating neurons

Fluoro-Jade C (Histo-Chem Inc.) stain was used to visualize degenerating neurons according to the manufacturer’s protocol in the target brain regions of ACC, amygdala, and PAG. There was no observable signal of neurodegeneration in the studied regions.

### Imaging and cell counting

Each histological section was digitized at 0.65 μm resolution (10 × magnification) using a Zeiss Axioscan microscope slide scanner. Images were then split into a separate channel for each fluorophore. Cell detection and counting were done with an open source QuPath software 0.3.2.^67^ . As each fluorescence channel was analyzed separately, the loci of immunofluorescence that were counted do not necessarily correspond to unique cells, especially for microglia and GABA channels where the fluorescent signal was more diffuse in appearance. Cell segmentation in brain regions downstream to the lesion site was done using the random trees algorithm embedded into QuPath. Cell segmentation of the lesion site was done by custom-trained Cellpose models (doi: 10.1038/s41592-022-01663-4). Although the researcher performing cell counts was blinded to the marmosets’ identity, it was possible to identify the site of the lesion and determine the animals group membership. Raw cell counts were transformed into cell densities per mm2 to account for size differences in ROI areas.

## Statistical analysis

All analyses were performed using R 4.2.3 (https://www.R-project.org/). For vocalization analysis, recording sessions were averaged by week and unless otherwise noted the data are represented as the mean values with mean ± confidence interval. Recording sessions obtained before the surgery were grouped as postnatal week 2 (pre-surgery) and all the recordings after the surgery were binned into postnatal weeks 3 to 6 (post-surgery). Due to COVID-19 related restrictions, recordings for some infants could not be extended beyond postnatal week 3.

Packages in tidyverse library^68^ were used for data processing and analyses. Linear mixed effect models (LMM) were used to analyze postnatal datasets, and these models were fitted with the lmer() function in lme4 package^69^. For estimation of coefficients, the maximum likelihood method was used. Models were fitted with postnatal weeks and experimental groups as fixed factors. To account for inter-individual variability, each monkey was modeled as a random effect. For multisyllabic phee analysis, syllables were nested within monkeys’ random effect. Models with and without lesions were used to test the effect of ACC lesion and lesion effect is considered significant at an αof 0.05^70,71^. Model assumptions were tested using the check_model() function available in the performance package. Log transformation was performed on some dataset to meet LMM model assumptions. When normality assumption was violated, non-parametric test (Wilcoxon test) was also used. Graphs were created using the package ggplot2.

For analysis involving immunofluorescence there were some inhomogeneities in the spatial distribution of IHC signal. The raw immunopositive detections for specific markers were normalized by NeuN cell counts that were obtained in the same Qupath processing pipeline. The final measure for each antigen-specific immunopositivity count was the ratio computed from the number of antigen-positive detections divided by the sum of the antigen-positive detections and NeuN-positive cells. This transformation bounded the possible antigen-specific detections between 0 and 1 and allowed for parametric modeling with beta distribution. Cell counts were fitted using glmmTMB software package in R statistical computing environment. As there were 3 sections per animal for each region of interest, these were modeled as random effects. Widths of major fiber tracts were modeled by a linear regression (function lm in base R). Final p values were adjusted for multiple comparisons with the false discovery rate method for each antigen.

## Acknowledgements

This research was supported by the Intramural Research Program of the National Institute of Mental Health (ZIAMH002951 and ZICMH002952 to YC). We thank George Dold, William Bennett, and David Ide from the NIMH Section on Instrumentation for customization of the stereotaxic frame and surgical anesthesia gas mask. We would also like to thank the Veterinary Medicine and Resources Branch and Central Animal Facility for animal husbandry, technical, and anesthetic support during procedures. GN is now at The Henry Jackson Foundation for the Advancement of Military Medicine, Bethesda, MD, USA. AP is now at Georgetown University, Washington DC, USA.

## Author Contributions (CRediT)

Conceptualization and methodology GN, YC; Software GN, DM, ACP, SPB; Formal analysis GN, DM; Investigation GN, DM, ACP; Resources YC; visualization GN, DM, YC; writing original draft GN, YC; Writing review and editing GN, DM, ACP, SPB, YC; Supervision DM, YC.

## Data Availability

All data needed to evaluate the conclusions in the paper are present in the paper.

---

**Fig. S1.**
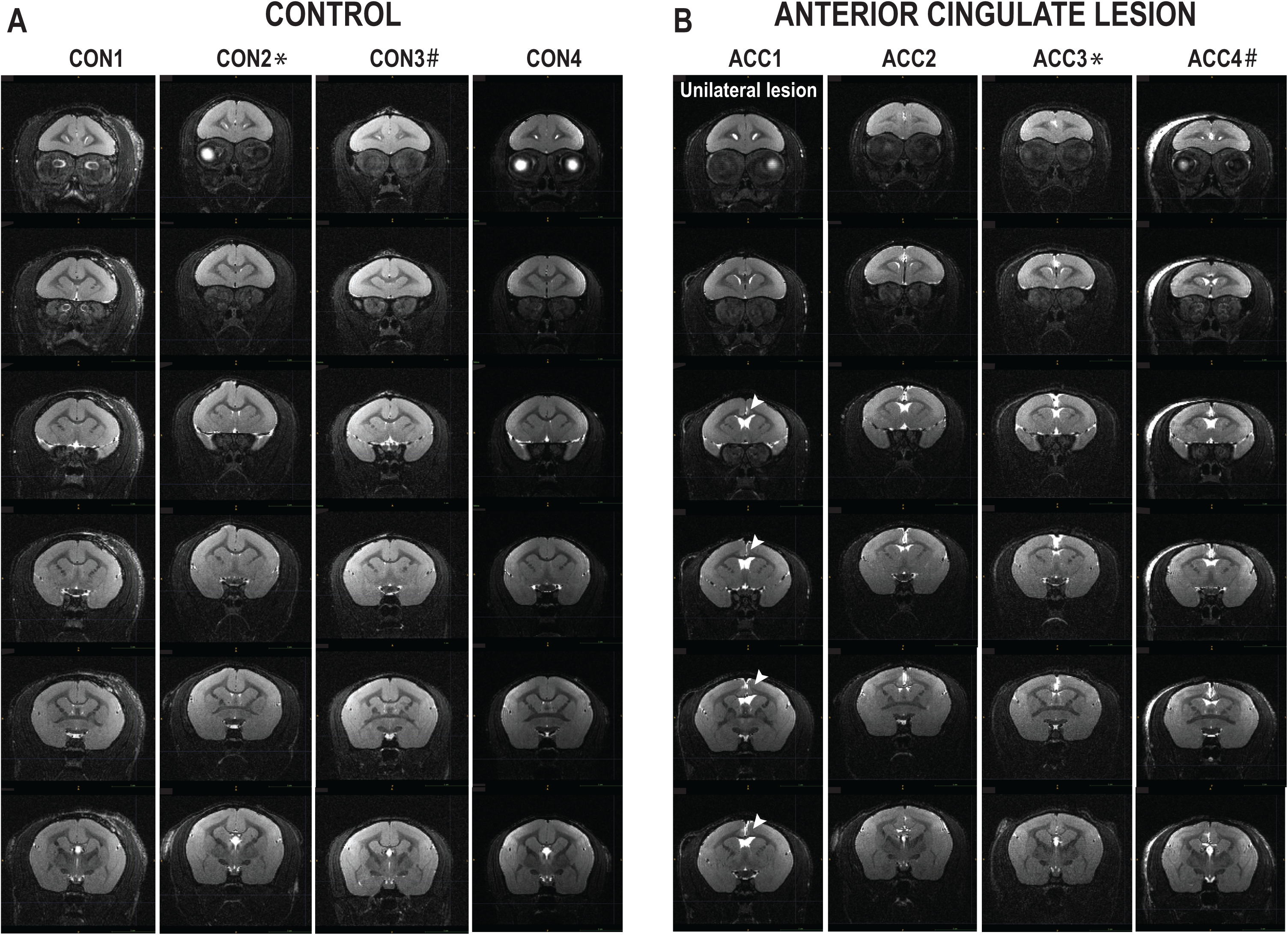
Bilateral MRI scans. Images from MRI scans of each marmoset representing coronal planes ranging from pregeniculate region until retrosplenial area. Each slice is 0.25mm apart. ACC1 shows a unilateral lesion depicted by the arrow. CON2* and ACC3* are twins, and CON3# and ACC4# are twins.

**Fig. S2.**
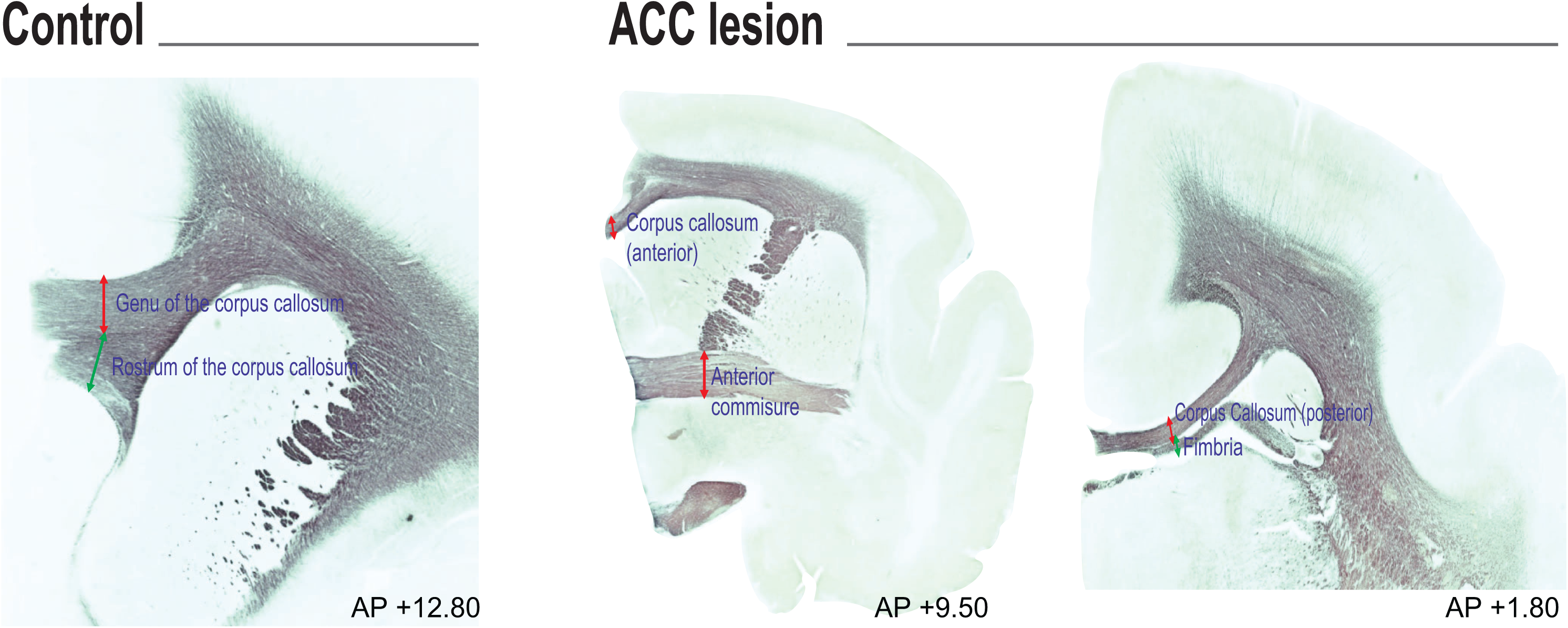
Representative images of white matter tract from a CON and ACC-lesioned subject. Both anterior and posterior corpus callosum are visibly narrowed in the area of the ACC lesion. The transverse widths of the white matter tracts were measured from sections stained for myelin at the following approximate rostrocaudal planes (in reference to the interaural axis): genu of the corpus callosum and rostrum of the corpus callosum (both +12.80 mm AP) were measured at the nadir of the overlaying cortex; anterior corpus callosum was measured at the midline and the anterior commissure was measured at the medial juncture with the internal capsule (both +9.50 mm AP); posterior corpus callosum and fimbriae were both measured at the junction with each other (both +1.80 mm AP).

**Fig S3.**
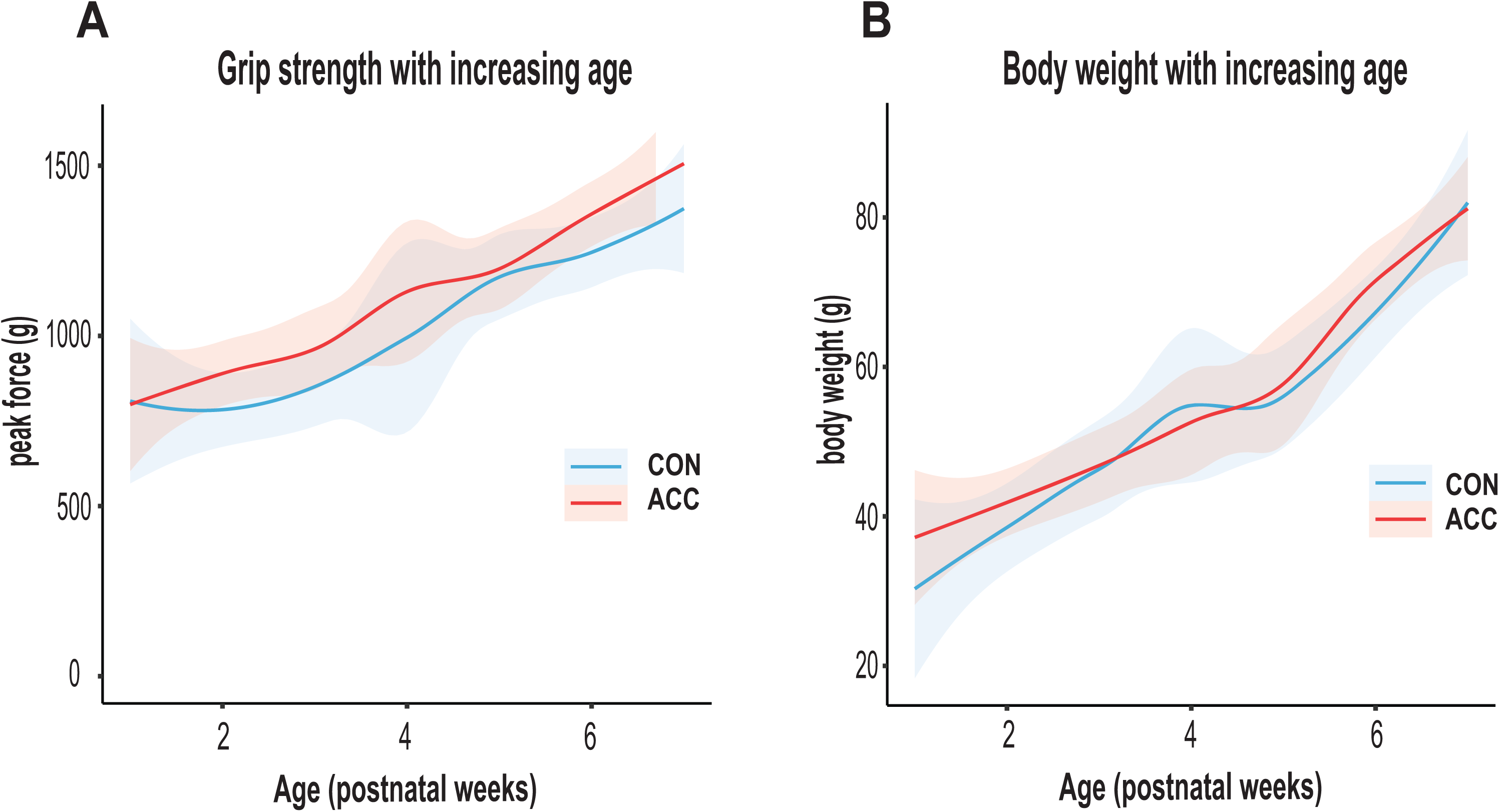
Physical factors in developing marmosets. **(A)** Muscular strength was quantified using the Bioseb (BIO-GS4) grip strength monitor. The infant was allowed to grip onto a bar while being gently pulled backwards in a horizontal plan to determine the maximal peak force (grams). The ACC lesion did not cause any changes to forelimb muscular strength. **(B)** Average body weight (grams) with increasing age. Animals in both groups showed comparable body weights with increasing age. CON (n=5), ACC (n=5).

